# Do GLP-1 Receptor Agonists Alter Brain Responses to Reward-Related Cues? A Systematic Review

**DOI:** 10.64898/2026.01.31.702984

**Authors:** Vincent Dang, Nicola Sambuco, Luba Yammine, Francesco Versace

**Author notes:** **Correspondence to:** Nicola Sambuco, Ph.D., Department of Translational Biomedicine and NeurosciencevUniversity of Bari Aldo MorovPiazza Giulio Cesare, 11, 70124 Bari, Italy. E-Mail, Francesco Versace, Ph.D., Department of Behavioral Science, Unit 1330, The University of Texas M. D. Anderson Cancer Center, PO Box 301439, Houston, TX 77230-1439. These authors contributed equally to this work (Vincent Dang, Nicola Sambuco).

## Abstract

Glucagon-like peptide-1 receptor agonists (GLP-1 RAs) are approved for treating type 2 diabetes and obesity and are under investigation as potential treatments for substance use disorders (SUD). GLP-1 RA-induced weight loss is thought to arise from both peripheral effects on gastrointestinal function and central modulation of appetite and reward circuits, though the exact mechanisms are unclear. Functional magnetic resonance imaging (fMRI) studies examining brain responses to reward-related cues can help clarify the central mechanisms through which GLP-1 RAs influence reward-seeking behavior. We systematically reviewed the fMRI literature examining how GLP-1 RAs affect brain responses to reward-related cues. We identified 1,209 records through a comprehensive literature search. After screening, only 11 studies met eligibility criteria. The vast majority assessed reactivity to food-related cues, with only one examining drug-related cues (alcohol), leaving neural mechanisms relevant to SUD largely unexplored. None included non-food emotional stimuli as control conditions. Several methodological limitations emerged. Most studies enrolled 20 or fewer participants per group, limiting statistical power. Treatment protocols varied substantially, with some assessing cue responses after single-dose administration and others after chronic treatment. Heterogeneity in medications used further confounds interpretation. The limited evidence tentatively suggests that acute GLP-1 RA administration may reduce brain reactivity to food cues in appetite and reward regions. However, effects appear inconsistent and may attenuate over time. Future studies should recruit larger samples, standardize agents and dosing, and assess responses to diverse motivationally relevant stimuli.

## Introduction

Glucagon-like peptide-1 receptor agonists (GLP-1 RAs) are a class of medications approved for the treatment of metabolic disorders such as type 2 diabetes mellitus (T2DM) and obesity (Wong et al., 2025). GLP-1 RAs mimic the effects of GLP-1, an endogenous hormone that increases insulin secretion, regulates blood glucose levels, delays gastric emptying, and promotes satiety (Friedman, 2024). Beyond improving glycemic control, GLP-1 RAs induce clinically meaningful weight loss in both people with and without diabetes (Davies et al., 2015; O’Neil et al., 2018; Wilding et al., 2021). The weight loss produced by GLP-1 RAs was initially attributed to their impact on the hypothalamic homeostatic circuits regulating satiety and appetite (Van Bloemendaal et al., 2014), but subsequent evidence from both animal and human studies showing that GLP-1 RAs reduce preference for energy-dense, palatable foods and blunt craving suggests that these medications might help reduce food intake by also modulating the hedonic aspects of eating (Blundell et al., 2017; Friedrichsen et al., 2021; Gabery et al., 2020). Recent retrospective and prospective studies in humans showed that GLP-1 RAs can influence reward-related behaviors beyond eating: use of GLP-1 RAs has been associated with changes in intake patterns of alcohol, nicotine, and other substances of abuse (Hendershot et al., 2025; Probst et al., 2023; Wang et al., 2024; Yammine et al., 2021). These cross-domain findings support the hypothesis that GLP-1 RAs broadly affect reward-related processes and underscore the need to elucidate the neurobiological mechanisms underlying these effects, which remain poorly understood.

The presence of GLP-1 receptors in brain regions central to reward processing (López-Ojeda & Hurley, 2024), together with evidence that GLP-1 RAs can modulate activity within these circuits (Gabery et al., 2020), has led to the hypothesis that GLP-1 RAs may reduce vulnerability to multiple maladaptive reward-seeking behaviors by blunting neuroaffective responses to reward-related cues (Eren-Yazicioglu et al., 2021; Krupa, 2025). Specifically, GLP-1 receptors are enriched in the lateral septum (where expression is highest), hypothalamus, amygdala, bed nucleus of the stria terminalis, hippocampus, ventral tegmental area (VTA), and nucleus accumbens (NAc). Critically, GLP-1-producing neurons in the nucleus tractus solitarius project monosynaptically to the VTA, NAc core, and NAc shell, projections that are activated by food intake, providing a direct anatomical substrate for GLP-1 modulation of reward processing. At the neurochemical level, preclinical studies demonstrate that GLP-1 RAs modulate dopamine levels and glutamatergic neurotransmission in reward circuits, attenuate accumbal dopamine release in response to drugs of abuse and suppress phasic dopamine responses to food-predictive cues in the VTA. These dopaminergic effects provide a specific neurochemical mechanism through which GLP-1 RAs may blunt cue-evoked motivational responses (Eren-Yazicioglu et al., 2021; Graham et al., 2020; Marquez-Meneses et al., 2025; Ong et al., 2017). This hypothesis is consistent with neurobiological models proposing that excessive eating and substance use disorders stem from a common mechanism: the attribution of excessive incentive salience to reward-related cues (Berridge et al., 2010; Robinson & Berridge, 2025; Stice & Yokum, 2016; Volkow et al., 2013). Incentive salience refers to the motivational properties of rewards and the cues associated with them. Stimuli with high incentive salience, such as hyperpalatable foods, capture attention, activate affective states, and motivate behaviors (Berridge, 2012). Through Pavlovian conditioning, cues that are repeatedly paired with consumption of hyperpalatable foods (e.g., the stretch of melted cheese on a slice of pizza in a commercial, the logo of a fast-food chain, or a dessert image on a menu) can acquire high incentive salience. Once cues hold high incentive salience, they become attractive, evoke intense craving, and trigger compulsive eating. In line with this theoretical model, neuroimaging studies show that food-related cues prompt stronger neuroaffective responses than neutral images, and these responses tend to be larger in individuals with obesity (Devoto et al., 2018; Hendrikse et al., 2015; Li et al., 2023).

Despite the theoretical rationale for hypothesizing that GLP-1 RAs affect neuroaffective responses to reward-related cues, empirical evidence is scarce. Anecdotal reports suggest that GLP-1 RAs might reduce “food noise”, a colloquial term used to describe persistent, intrusive thoughts about food that is conceptually related to the well-established construct of craving (Dhurandhar et al., 2025). Clinical trials also report decreases in self-reported food cravings (Blundell et al., 2017; Friedrichsen et al., 2021). However, subjective measures do not provide direct evidence that GLP-1 RAs reduce neurophysiological responses to food-related or other reward-related cues. Determining the extent to which GLP-1 RAs modulate neuroaffective responses to food-related cues and other motivationally relevant stimuli is critical both for understanding their mechanisms of action and for assessing their promise as treatments for maladaptive reward-seeking behaviors, including substance use disorders.

Although recent reviews have examined GLP-1 signaling and reward processing (Au et al., 2025; Badulescu et al., 2024; Schulz et al., 2023), none has focused specifically on cue-induced brain responses measured with task-based fMRI, the neural signals most relevant to understanding how GLP-1 RAs might reduce the motivational pull of rewarding stimuli, or critically evaluated the methodological approaches in this emerging literature. This review aimed to systematically evaluate the neuroimaging literature on the effects of GLP-1 RAs on reactivity to reward-related cues, whether food-related cues in individuals with obesity or drug-related cues in individuals with substance use disorders. A secondary objective was to evaluate the methodological approaches used in these studies to identify best practices for investigating and interpreting the effects of GLP-1 RAs on neuroaffective responses to motivationally relevant stimuli.

## Method

### Inclusion/Exclusion Criteria

Inclusion criteria included studies with human subjects over the age of 18 years who received any GLP-1 RA (e.g., exenatide, semaglutide) as the intervention. Because this review focused on brain responses to reward-related cues, eligible studies had to assess reactivity to food-related, or drug-related cues using fMRI. Exclusion criteria included non-human studies, articles not in English, and review articles (e.g., systematic reviews, meta-analyses).

### Information Sources

The literature search for all databases was conducted in January 2026. Studies were pulled from Embase, Web of Science, Scopus, Medline, PubMed, the Cochrane Library, PsycINFO, and Google Scholar. The search string for each database was approved and finalized by an MD Anderson Librarian. An example search string used for Embase is: GLP-1.mp. or glucagon like peptide 1/ AND reward/ or monetary reward/ or (“food cue*” or addiction* or cocaine or alcohol or smoking or “substance use” or reward* or “substance abuse” or marijuana or meth).mp AND stimulus/ or (stimul* or brain response*).mp. The search strings used for the other databases can be found in the supplemental information. Grey literature sources and clinical trial registries were also searched to identify ongoing or unpublished studies.

## Study Selection

The study protocol was preregistered on OSF on December 13, 2024 (10.17605/OSF.IO/JBPV7). This systematic review followed the 2020 Preferred Reporting Items for Systematic Review and Meta-Analyses (PRISMA) guidelines (Page et al., 2021). All study references were exported to EndNote and imported to the Covidence systematic review management platform. Covidence was used to remove duplicate references pulled from the databases. The first author (VD) independently screened through the titles and abstracts and brought up any eligibility concerns with the last author (FV). After finalizing all abstracts with the last author, the first author conducted a full-text review on the selected texts. All questions and eligibility concerns were discussed and finalized with both authors. Data extracted included the study title, study design, sample size, study population, GLP-1RA type, dose, duration, route of administration, type of measure used to record brain responses, and main outcome of each study. The main outcome of interest was effect of GLP-1RAs on cue reactivity.

## Results

### Primary Search Results

The literature search identified a total of 1,209 studies across all databases. After removing 368 duplicate studies using the Covidence software and an additional 162 duplicates manually, 679 studies were screened for eligibility based on title and abstract. Of these, 49 were deemed eligible for full-text screening. After full-text review, 38 studies were excluded, resulting in 11 studies being included in the final data extraction (see **Figure 1** for the PRISMA flow diagram). Despite the widespread clinical use and scientific interest surrounding GLP-1 RAs, only a handful of neuroimaging studies have investigated their effects on brain activity, and these studies are methodologically heterogeneous.

**Figure 1.**
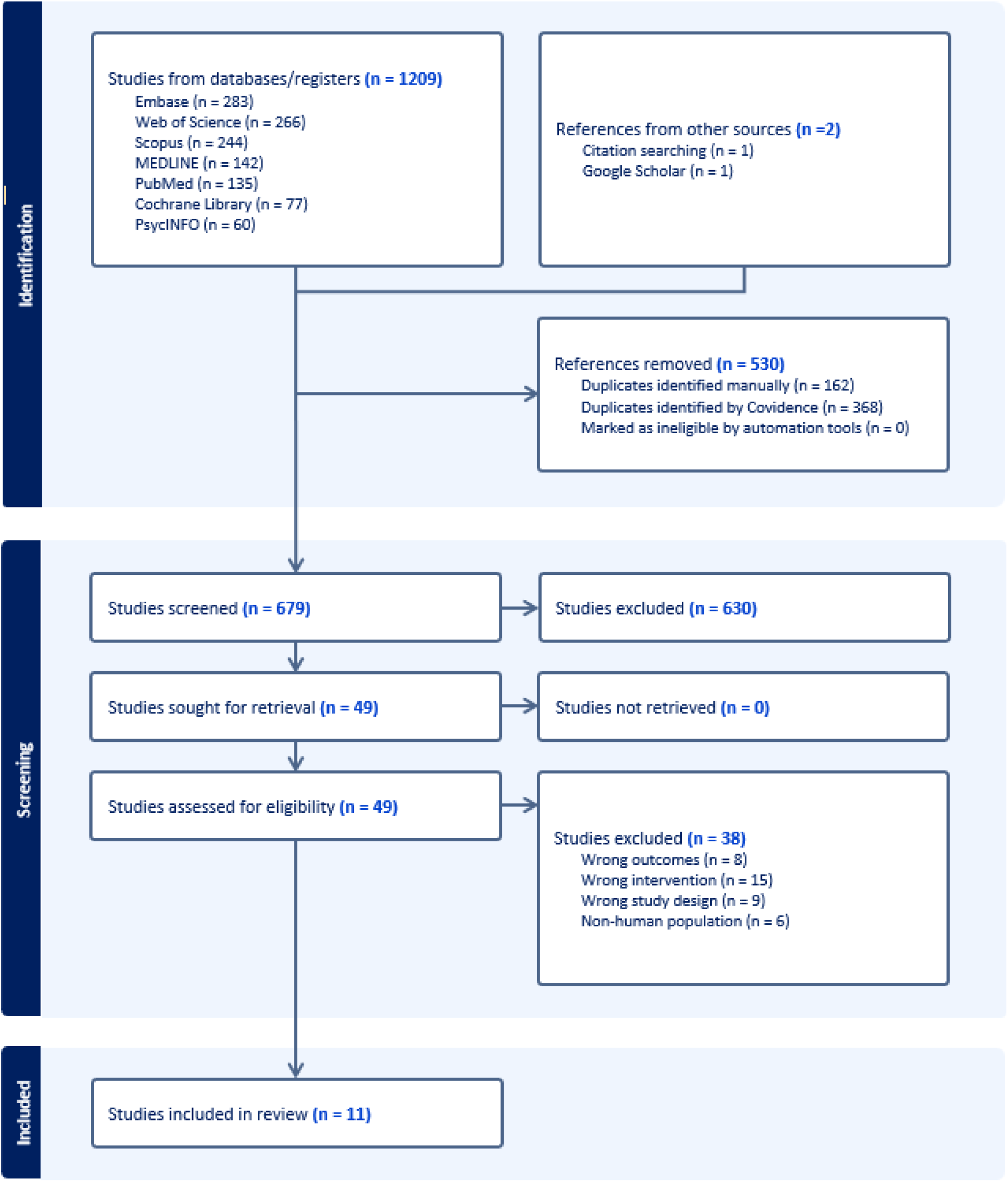
PRISMA flow diagram of the study selection process. A supplementary search of grey literature sources and clinical trial registries did not identify additional published studies.

### Overview of the Neuroimaging Evidence Base

Across the 11 studies identified, six used infusions to examine the acute effects of GLP-1 RAs on cue reactivity, and five assessed long-term (5 weeks to 26 weeks) treatment effects. Sample sizes were generally small, with 10 out 11 studies including fewer than 20 participants per group. Most studies included only neutral and food-related cues (ten studies); only one included alcohol-related cues (but not food-related cues), and none included non-food-related motivationally relevant cues as active control conditions to evaluate the selectivity of GLP-1 RA effects. Eight studies analyzed brain activity only within selected regions of interest (ROI). Although this approach might have been driven by the small sample sizes, the ROIs were heterogeneous across studies, and no clear activation patterns emerged. Most studies used exenatide (n = 5) or liraglutide (n = 3), with single studies examining lixisenatide or GLP-1 7-36 amide. Only one study (Martin et al., 2025) tested a newer-generation agent, tirzepatide, a dual GIP/GLP-1 receptor agonist.

In summary, this limited and methodologically heterogeneous evidence base constrains the conclusions that can be drawn about how GLP-1 RAs affect brain responses to food-or drug-related cues, and substantial gaps remain in our understanding of their effects on central mechanisms of motivation and reward. **Table 1** summarizes the characteristics and main findings of the 11 included studies. In the sections that follow, we examine these findings in greater detail.

**Table 1.**
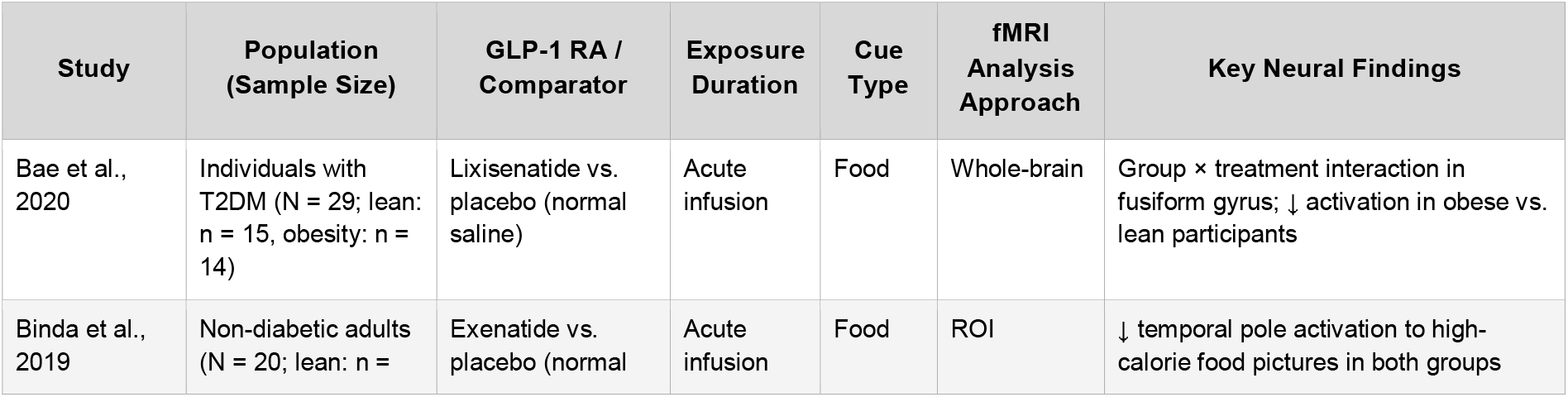
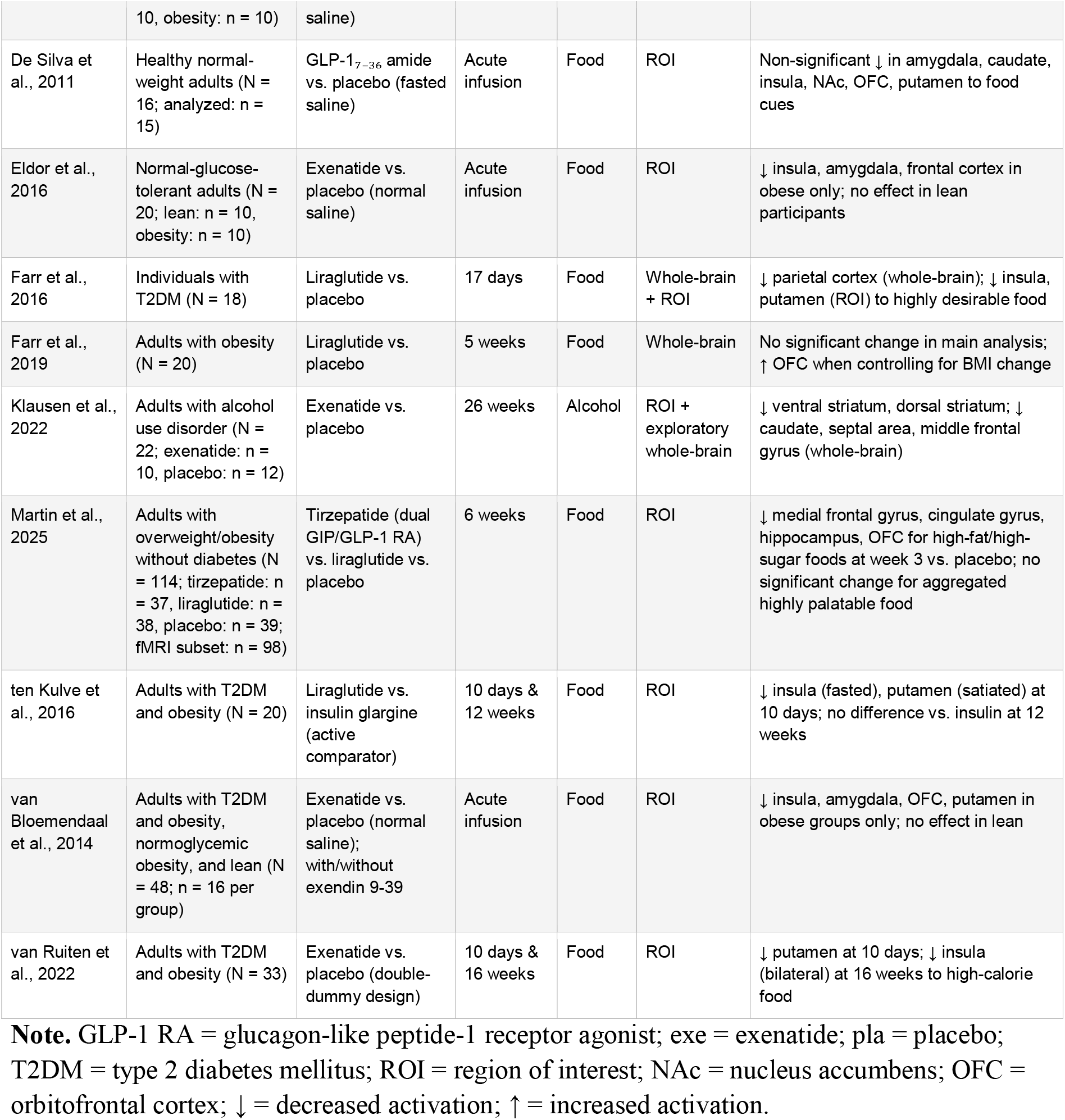
Summary of Studies Included in the Review.

The included studies exhibited considerable methodological variability. **Table 2** summarizes the key methodological characteristics observed across these studies. All studies except one included fewer than 20 participants per group. Studies varied in treatment duration (acute infusion to 26 weeks), GLP-1 RA agents used (reflecting the medications available at the time of each study), stimulus categories examined, and analytic approaches employed. This heterogeneity limits cross-study comparisons.

**Table 2.** Summary of Methodological Limitations Across Included Studies.

| Methodological Considerations | Description |
| --- | --- |
| Small sample sizes | All studies except one included <20 participants per group, limiting statistical power for between-group comparisons |
| Acute vs. long-term effects | 6 studies examined acute effects; 5 examined long-term (5-26 weeks), complicating cross-study comparisons |
| Medication heterogeneity | Exenatide (5 studies), liraglutide (3 studies), lixisenatide (1), GLP-1 7-36 amide (1), tirzepatide (1); only one study tested a newer-generation agent (tirzepatide, a dual GIP/GLP-1 RA) |
| Limited cue types | 10 studies used food cues only; 1 used alcohol cues; none used non-food/non-drug emotional stimuli as control |
| Analysis approach | 8 studies used ROI analyses only, with heterogeneous ROI selection; 3 used whole-brain analyses |

### Regional Effects of GLP-1 RAs on Brain Cue Reactivity

Across the eleven studies included in this review, GLP-1 RA administration was associated with altered neural responses in eleven brain regions during food cue processing (**Figure 2; Table 3**). The upper panel of **Figure 2** illustrates the direction of these effects: blue regions showed predominantly decreased BOLD responses, red regions showed increased responses, and yellow regions exhibited mixed findings across studies. The lower panel displays the number of studies reporting effects in each region, with darker shading indicating higher convergence.

**Table 3.**
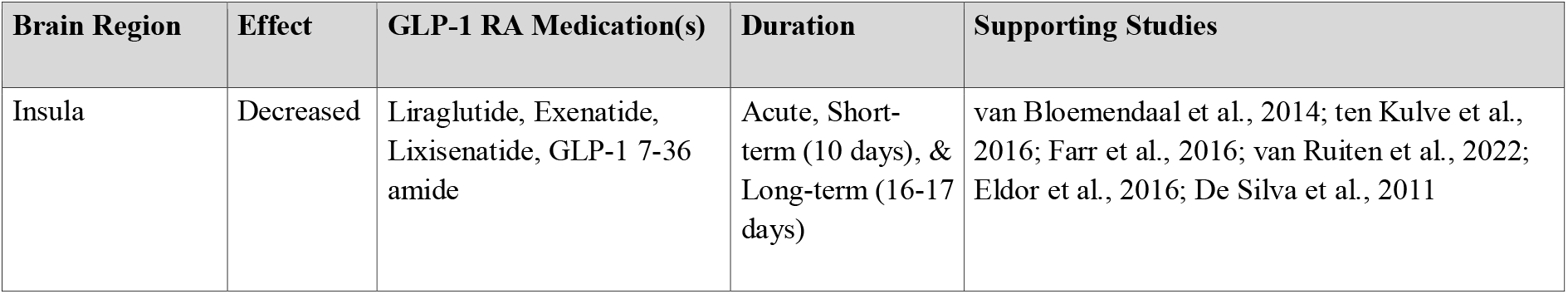
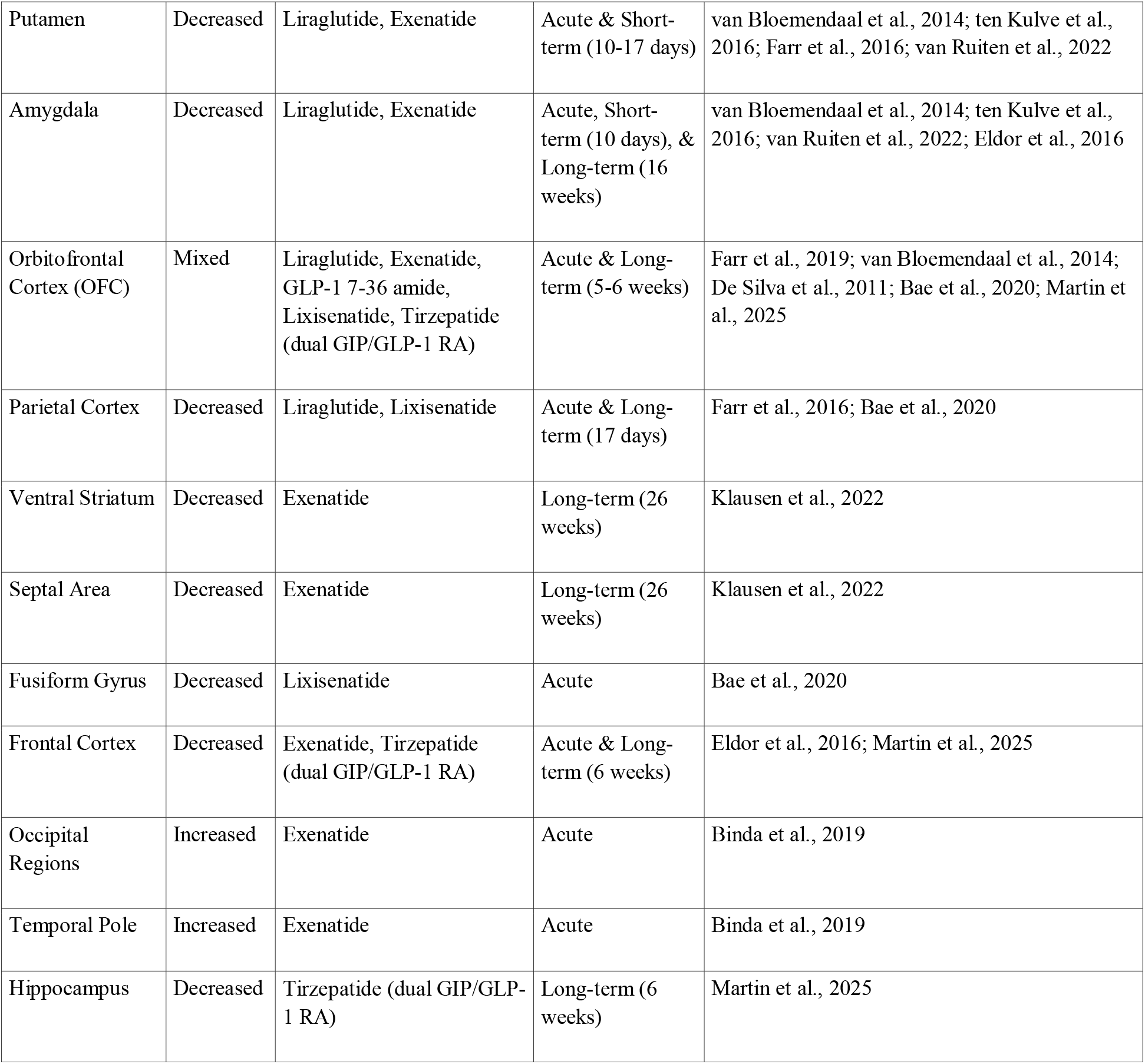
Summary of brain regions showing altered cue reactivity following GLP-1 RA administration.

| Brain Region | Effect | GLP-1 RA Medication(s) | Duration | Supporting Studies |
| --- | --- | --- | --- | --- |
| Insula | Decreased | Liraglutide, Exenatide, Lixisenatide, GLP-1 7-36 amide | Acute, Short-term (10 days), & Long-term (16-17 days) | van Bloemendaal et al., 2014; ten Kulve et al., 2016; Farr et al., 2016; van Ruiten et al., 2022; Eldor et al., 2016; De Silva et al., 2011 |
| Putamen | Decreased | Liraglutide, Exenatide | Acute & Short-term (10-17 days) | van Bloemendaal et al., 2014; ten Kulve et al., 2016; Farr et al., 2016; van Ruiten et al., 2022 |
| Amygdala | Decreased | Liraglutide, Exenatide | Acute, Short-term (10 days), & Long-term (16 weeks) | van Bloemendaal et al., 2014; ten Kulve et al., 2016; van Ruiten et al., 2022; Eldor et al., 2016 |
| Orbitofrontal Cortex (OFC) | Mixed | Liraglutide, Exenatide, GLP-1 7-36 amide, Lixisenatide, Tirzepatide (dual GIP/GLP-1 RA) | Acute & Long-term (5-6 weeks) | Farr et al., 2019; van Bloemendaal et al., 2014; De Silva et al., 2011; Bae et al., 2020; Martin et al., 2025 |
| Parietal Cortex | Decreased | Liraglutide, Lixisenatide | Acute & Long-term (17 days) | Farr et al., 2016; Bae et al., 2020 |
| Ventral Striatum | Decreased | Exenatide | Long-term (26 weeks) | Klausen et al., 2022 |
| Septal Area | Decreased | Exenatide | Long-term (26 weeks) | Klausen et al., 2022 |
| Fusiform Gyrus | Decreased | Lixisenatide | Acute | Bae et al., 2020 |
| Frontal Cortex | Decreased | Exenatide, Tirzepatide (dual GIP/GLP-1 RA) | Acute & Long-term (6 weeks) | Eldor et al., 2016; Martin et al., 2025 |
| Occipital Regions | Increased | Exenatide | Acute | Binda et al., 2019 |
| Temporal Pole | Increased | Exenatide | Acute | Binda et al., 2019 |
| Hippocampus | Decreased | Tirzepatide (dual GIP/GLP-1 RA) | Long-term (6 weeks) | Martin et al., 2025 |

**Figure 2.**
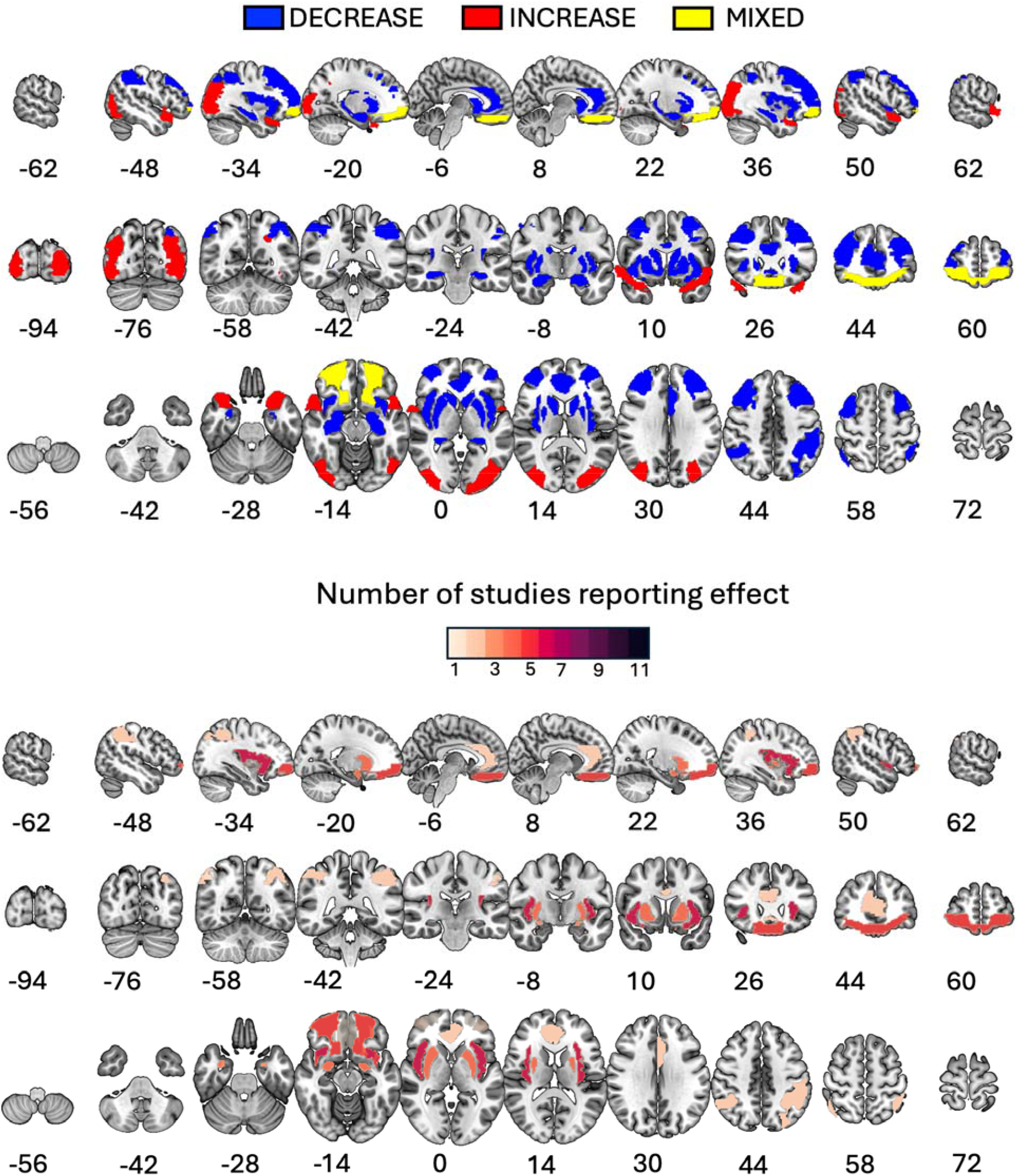
Summary of brain regions showing changes in cue-evoked activation (i.e., cue reactivity) following GLP-1 RA administration across eleven studies. The upper panel displays the direction of effects: regions showing predominantly decreased BOLD responses (blue), increased responses (red), or mixed findings across studies (yellow). The lower panel displays the number of studies (1–11) reporting effects in each region, with darker shading indicating greater convergence across studies. Slices are displayed in MNI space; coordinates indicate slice position in mm.

The most consistent finding was reduced activation, with eight regions showing predominantly decreased BOLD responses following GLP-1 RA treatment. These included the insula, putamen, amygdala, parietal cortex, ventral striatum, septal area, fusiform gyrus, and frontal cortex. In contrast, the occipital cortex and temporal pole showed increased activation (Binda et al., 2019). Findings in the orbitofrontal cortex were mixed, with some studies reporting increases and others reporting decreases.

The proportion of studies reporting effects varied substantially across regions (**Figure 2**, lower panel). The insula emerged as the most consistently implicated region, with six out of eleven studies reporting GLP-1 RA-related changes in this area. The putamen and amygdala were each implicated in four studies, and OFC in five. Notably, several regions were reported in only a single study, including the ventral striatum, septal area, fusiform gyrus, frontal cortex, occipital regions, hippocampus, and temporal pole. The uneven cross-study convergence should be considered when interpreting the robustness of region-specific findings. **Table 3** summarizes the regional findings, including the direction of effects, medications used, administration duration, subject populations, and supporting studies.

#### Whole-Brain Analyses of GLP-1 RA Effects on Food Cue Reactivity

Whole-brain analyses yielded inconsistent results. Acute lixisenatide administration produced opposite effects in lean and obese participants viewing food versus non-food images, with a significant group-by-treatment interaction in the fusiform gyrus (Bae et al., 2020). The two liraglutide studies did not replicate these effects. After 17 days of treatment, liraglutide reduced activation in the inferior parietal cortex in response to highly desirable versus less desirable food images (Farr et al., 2016), but after 5 weeks of treatment, no significant differences emerged in the main analysis (Farr et al., 2019). Both liraglutide studies reported results from secondary ROI analyses: Farr et al. (2016) found reduced activation in the insula and putamen, while Farr et al. (Farr et al., 2019) found increased orbitofrontal cortex activation when controlling for BMI/weight changes.

In summary, whole-brain analyses do not reveal a consistent pattern of GLP-1 RA effects on cue-elicited activity. This lack of convergence likely reflects methodological heterogeneity and limited statistical power rather than true null effects.

#### Region-of-Interest Analyses of GLP-1 RA Effects on Food Cue Reactivity

Seven studies employed ROI-based analyses. The ROIs chosen converged on key nodes in the appetitive and reward network, most consistently including the amygdala (5 studies), insula (6 studies), striatum (putamen: 5 studies; caudate: 3 studies; nucleus accumbens: 2 studies), and orbitofrontal cortex (5 studies). Only one study included the hippocampus and hypothalamus, and one secondary analysis focused on occipital visual areas.

The results were mixed both within and across studies, with no clear pattern emerging in any ROI examined by more than one study, except perhaps in the insula, where the majority of studies reported decreased activation to food-related cues after short-or long-term GLP-1 RA administration (see **Figure 2**). Compared with insulin glargine, 10 days of liraglutide treatment decreased activation to high-calorie food pictures in the insula (bilaterally) of participants with type 2 diabetes, but only after overnight fasting; this effect was no longer present after 12 weeks of treatment (ten Kulve et al., 2016). Secondary analyses by Farr et al. (2016) similarly showed that acute liraglutide administration decreases insula activation in fasted participants with T2DM. Acute exenatide administration also decreased activation in the insula in obese participants with and without T2DM, but not in lean participants (Eldor et al., 2016; Van Bloemendaal et al., 2014). Unlike ten Kulve et al. (2016), Van Ruiten et al. (2022) found no effect after 10 days of treatment, but reported that prolonged (16 weeks) treatment with exenatide decreased activation in response to food pictures in the insula. Finally, De Silva et al. (2011) reported a non-significant trend toward reduced activity in the right insula when viewing food pictures after infusion of GLP-□□□□ amide.

Activity in all other ROIs showed inconsistent responses to GLP-1 RAs, with approximately half of the studies showing decreased activation in response to food-related cues and the other half showing no changes (see **Figure 2**, bottom panel).

### GLP-1 RA Effects on Alcohol Cue Reactivity

Only one study examined GLP-1 RA effects on drug-related cue reactivity. At 26 weeks, Klausen et al. (2022) found that participants treated with exenatide exhibited significantly lower alcohol cue reactivity in the ventral striatum compared to the placebo group. No significant group differences were detected in the dorsal striatum or putamen, the other ROIs analyzed. When assessing within-group changes, exenatide significantly reduced alcohol cue reactivity from baseline to week 26 in both the ventral striatum and the dorsal striatum, but not in the putamen. No significant changes were observed in the placebo group across the 26-week period. An exploratory whole-brain analysis further indicated that exenatide significantly reduced alcohol cue reactivity in the left caudate, septal area, and right middle frontal gyrus at the 26-week time point.

Regarding clinical outcomes, exenatide did not significantly reduce the number of heavy drinking days compared with placebo in the overall sample. However, a significant reduction in both heavy drinking days and total alcohol intake was observed in the subset of participants classified as obese (BMI >30 kg/m2).

### Supplementary Search: Ongoing Studies

Given the growing interest in repurposing GLP-1 medications for substance use disorders (Yammine et al., 2025), several ongoing and planned study protocols are implementing neuroimaging tasks to rigorously examine the mechanism of GLP-1 in reducing brain reactivity to drug-related cues (Freet et al., 2024; Klausen et al., 2025; Simmons, 2023; Yammine et al., 2023). These double-blinded randomized controlled trials may help add methodologically robust studies to the GLP-1 and neuroimaging literature. A supplementary search also identified Atila et al. (2022), who found no significant difference in whole-brain functional activity between dulaglutide and placebo groups in response to smoking cue videos after 12 weeks of treatment.

## Discussion

Functional neuroimaging studies investigating the effects of GLP-1 receptor agonists on brain responses to reward-related cues are scarce and methodologically heterogeneous. Given the growing therapeutic relevance and potential widespread use of these medications, this gap in understanding is concerning, as the neural effects of GLP-1 RAs on the reward system remain poorly characterized and their mechanisms of action not completely understood.

Most available studies have focused on reactivity to food-related cues, with only one examining drug-related cues and none exploring reactivity to other motivationally relevant or emotional stimuli. This imbalance is understandable, as GLP-1 RAs were originally developed for diabetes and weight management, and their effects on substance use have been discovered more recently. Nevertheless, as several clinical trials are now underway to test GLP-1 RAs in substance use disorders, it is critical to characterize their effects on the reward system to understand the mechanisms these agents may engage to reduce drug-related motivation.

Even within the food domain, several methodological issues make summarizing data across studies difficult: studies enrolled small sample sizes, used diverse pharmacological agents (e.g., exenatide, liraglutide, lixisenatide) with different dosing regimens, and mixed acute versus chronic designs. Despite these limitations, a tentative pattern emerges: acute GLP-1 RA administration might reduce brain reactivity to high-calorie food cues within key reward and salience regions, including the striatum, anterior cingulate cortex (ACC), insula, and orbitofrontal cortex (OFC). However, these effects are inconsistent across studies and often depend on analytic strategies; most positive findings derive from region-of-interest (ROI) analyses, which may inflate apparent significance and bias interpretation (Gentili et al., 2019). In contrast, studies of long-term administration report weaker or absent modulation of these circuits, suggesting that neural adaptations may normalize with sustained exposure or that acute effects reflect transient pharmacodynamic responses rather than stable neuroplastic changes.

Collectively, these preliminary results suggest that GLP-1 RAs might modulate brain activity in regions involved in both sensory appraisal and reward valuation, potentially reducing the motivational salience of food-related cues. However, because existing studies have compared only neutral and food-related stimuli, it remains unclear whether GLP-1 RAs selectively attenuate responses to food cues or instead produce a more generalized dampening of affective and motivational reactivity. To clarify the central mechanisms of GLP-1 action, future work must include larger samples, harmonize drug regimens, and assess brain responses to a wider range of appetitive and aversive stimuli. Incorporating whole-brain analyses and rigorous control conditions will be essential to disentangle whether the observed neural alterations are specific to reward processing or reflect broader effects on visual attention, salience attribution, or emotional regulation.

### Methodological considerations and limitations of the regional summary

The regional summary of neuroimaging findings (**Figure 2**) indicates that GLP-1 RA effects have been reported in brain areas consistently identified in basic affective neuroscience research as responsive to the emotional content of visual stimuli, including the insula, amygdala, and orbitofrontal cortex (Frank et al., 2019; Sabatinelli et al., 2006, 2007, 2011; Sambuco, Bradley, et al., 2020). However, several considerations temper the strength of conclusions that can be drawn from these findings. First, the proportion of studies reporting effects varied substantially across regions, with only the insula implicated in more than half of the studies reviewed (6 of 11), while several regions were reported in only a single study. Notably, the largest study to date (Martin et al., 2025)(N=114) found no significant effects in the insula with tirzepatide despite robust behavioral effects on food intake, suggesting that insula findings from smaller studies may not be reliable or that effects vary by medication class. Second, many of the included studies relied on region-of-interest (ROI) analyses, which, while offering increased sensitivity for detecting effects in a priori specified regions, may inflate the apparent consistency of findings when aggregated across studies. As Gentili and colleagues (Gentili et al., 2019) demonstrated in an activation likelihood estimation (ALE) meta-analysis of specific phobia, including ROI-based results can substantially change meta-analytic conclusions relative to analyses restricted to whole-brain voxel-wise approaches alone. In their analyses, some regions (e.g., amygdala, insula) appeared robustly implicated when ROI studies were included but did not survive when analyses were limited to whole-brain findings. Given that the present review similarly aggregates across ROI and whole-brain studies, the regional convergence observed here should be interpreted with caution. Future research employing larger samples and whole-brain analytic approaches will be essential to establish a more definitive map of brain regions modulated by GLP-1 RA administration during food cue processing.

A related consideration concerns the selection of ROIs in the reviewed studies. While many targeted regions plausibly involved in affective/reward processing, several selected areas lack strong support from the affective neuroscience literature that used pictures to evoke emotional responses. For instance, the hippocampus, examined in only one study, is not uniformly engaged during the presentation of emotional stimuli; the anterior hippocampus responds to affective visual content (Bradley & Sambuco, 2022; Sambuco & Bradley, 2024), but posterior regions do not (Bradley et al., 2022; Sambuco, Bradley, et al., 2020), meaning an ROI encompassing the entire structure may dilute detectable effects. Even the insula, the most frequently reported region in this review, may not be central to affective processes per se. Research over the past decade has challenged the notion of a single “core affect network” activated uniformly across emotional contexts (Lindquist et al., 2012), demonstrating instead, unsurprisingly, that neural responses depend heavily on induction modality and task demands (Sambuco, Bradley, et al., 2020; Sambuco, Costa, et al., 2020b; Sambuco, Bradley, et al., 2025). The insula, for example, is reliably activated during threat anticipation (Sambuco, Costa, et al., 2020b, 2020a) but less so during free viewing of emotional images, where inferior frontal gyrus (IFG) activation near opercular regions may be the primary driver (Sambuco, Bradley, et al., 2020), and spatial smoothing could misattribute these effects to the adjacent insula. Current evidence suggests emotional visual perception engages a distributed network spanning visual cortex, lateral posterior parietal regions, subcortical structures (amygdala, anterior hippocampus), lateral (particularly IFG) and medial prefrontal regions (Sabatinelli et al., 2011; Sambuco, 2022; Sambuco, Bradley, et al., 2020). Whether the insula activation observed here reflects food-specific processing or a broader affective response remains to be determined; future studies could disentangle these possibilities by comparing reactivity to food cues with reactivity to other affective, non-food-related stimuli within the same paradigm.

### Mechanistic Interpretation

From a neurobiological standpoint, these findings align with evidence that GLP-1 receptors are expressed throughout the brain, including the nucleus accumbens, ventral tegmental area, hypothalamus, and prefrontal cortex (Gupta et al., 2023). GLP-1 signaling modulates dopaminergic transmission within these networks, reducing phasic dopamine release in response to reward cues and thereby decreasing incentive salience attribution (Robinson & Berridge, 2025). The observed attenuation in striatal and ACC activity could therefore reflect reduced dopaminergic input and diminished engagement of the valuation network.

These data also align with the incentive sensitization framework, which posits that exaggerated motivational responses to reward-predictive cues, rather than enhanced hedonic pleasure, drive compulsive consumption (Robinson & Berridge, 2025). GLP-1-induced reductions in neural reactivity to food-related cues within valuation circuits may therefore counteract incentive sensitization mechanisms, providing a plausible account of reduced craving reported by patients. However, given the small number of available studies and the limited reproducibility of fMRI cue-reactivity paradigms with few trials and participants, such interpretations should be viewed as provisional.

An additional consideration is whether GLP-1 RA effects on brain activity depend on specific food stimulus categories. Martin et al. (2025) found that tirzepatide decreased activation in the medial frontal gyrus, cingulate gyrus, hippocampus, and orbitofrontal cortex specifically in response to high-fat, high-sugar food images, but not to the aggregated highly palatable food category. This pattern may reflect genuinely selective effects on neural responses to the most energy-dense foods, or it may arise because responses to other food categories were near baseline levels, effectively washing out effects in the combined analysis. Without direct comparisons of BOLD signal amplitude across food categories, definitive conclusions about stimulus specificity cannot be drawn (Bradley et al., 2017; Sambuco, Stevens, et al., 2025; Schupp et al., 2000; Weinberg & Hajcak, 2010).

### Future Research Directions

To move beyond preliminary observations, several key research questions must be addressed (see Table 4 for summary).

**Table 4.**
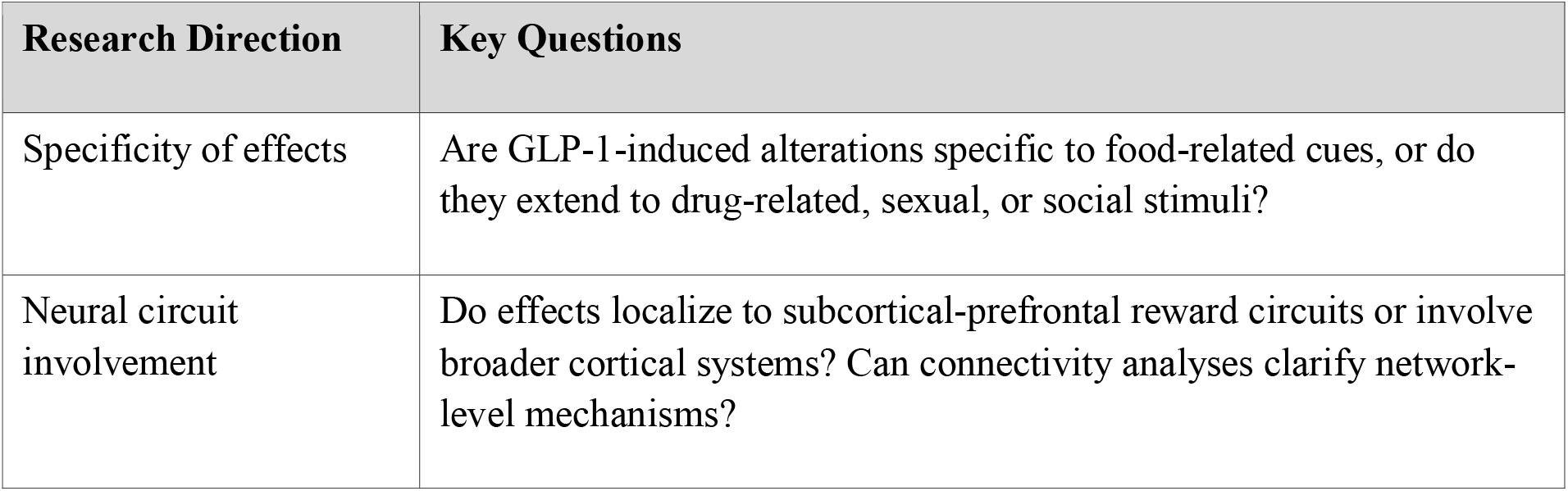
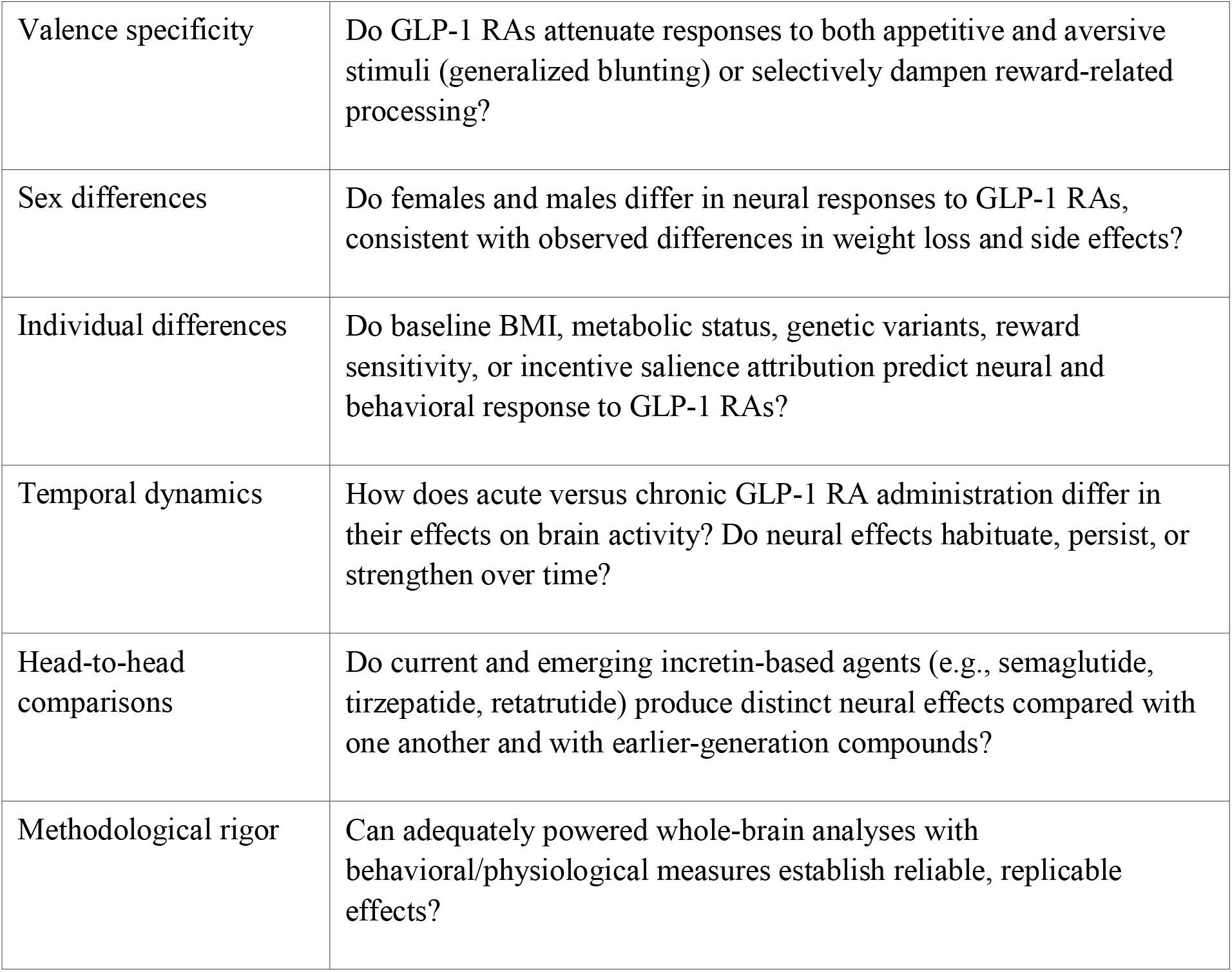
Future Research Directions and Key Questions.

#### Specificity of effects

Are GLP-1-induced alterations specific to food-related cues, or do they extend to other appetitive categories such as drug-related, sexual, or social stimuli? Addressing this question will clarify whether GLP-1 modulation targets domain-specific appetite regulation or exerts a broader influence on motivational salience.

#### Neural circuit involvement

Do these effects localize primarily to subcortical-prefrontal circuits traditionally associated with reward processing, or do they involve broader cortical systems encompassing parietal and visual regions typically engaged during the processing of motivationally relevant visual stimuli? Combining univariate fMRI analyses with connectivity analyses could determine whether GLP-1 primarily dampens striatal valuation processes or disrupts the integration of sensory and motivational information across distributed networks.

#### Valence specificity

If GLP-1 RAs attenuate neural responses to both appetitive and aversive stimuli, this pattern could indicate a generalized dampening of emotional reactivity. Notably, affective blunting has been reported as a side effect of other commonly prescribed medications, including SSRIs and beta blockers (MacCormack et al., 2021; Peters et al., 2022). Conversely, if GLP-1-related blunting is selective to reward-related content, these compounds may normalize aberrant hyperreactivity in individuals with obesity or substance use disorders without reducing negative affective processing. Carefully designed experiments that manipulate both valence and motivational intensity are essential.

#### Sex differences

Recent literature suggests that females tend to achieve greater weight loss and experience more gastrointestinal side effects than males (Yang et al., 2025; Marassi et al., 2025; Inceu et al., 2025; Rentzeperi et al., 2022). Future cue reactivity fMRI studies may benefit from examining sex as a potential moderating variable.

#### Individual differences

Beyond sex, baseline individual differences in the tendency to attribute incentive salience to cues may moderate neural responses to GLP-1 RAs. Individuals with heightened cue reactivity at baseline may show greater treatment-related reductions, or alternatively, may be more resistant to pharmacological modulation. Future studies should assess baseline reward sensitivity and incentive salience attribution as potential predictors of response.

#### Temporal dynamics

The existing literature includes both acute infusion and chronic treatment designs, yet direct comparisons of short-versus long-term effects within the same study are rare. Understanding whether neural effects emerge rapidly, habituate with continued exposure, or strengthen over time is critical for optimizing treatment protocols.

#### Head-to-head comparisons

The included studies used different medications with varying receptor selectivity (e.g., GLP-1 selective vs. dual GIP/GLP-1 agonists), half-lives, and CNS penetrance. Notably, several of the GLP-1 RAs examined in these studies (e.g., exenatide, lixisenatide) are gradually being retired from clinical use, while the therapeutic landscape is shifting toward newer GLP-1 RAs (e.g., semaglutide) and dual-or triple-agonist compounds (e.g., tirzepatide, retatrutide). Future neuroimaging studies should therefore prioritize head-to-head comparisons among current and emerging incretin-based agents to determine whether the heterogeneity observed across existing studies reflects true pharmacological differences, such as variation in receptor selectivity, CNS penetrance, or half-life, or methodological variability.

#### Methodological rigor

Studies should adopt adequately powered samples and prioritize whole-brain voxel-wise analyses to enable unbiased identification of GLP-1 RA effects across the brain. ROI analyses may complement whole-brain findings by providing increased sensitivity for hypothesis-driven questions, but results from these two approaches should be reported separately to facilitate future meta-analytic synthesis. Incorporating behavioral and physiological measures of motivation, such as cue-induced craving or attentional bias indices, will facilitate cross-modal validation of neuroimaging findings.

## Conclusions

Current evidence suggests that GLP-1 receptor agonists might modulate neural systems underlying reward processing by reducing activity within valuation and motivational circuits. These findings provide a plausible neurobiological basis for the reported behavioral effects of GLP-1 RAs on appetite, craving, and potentially substance use. Yet the field remains in its early stages. Establishing the reproducibility, specificity, and clinical significance of these neural changes will require standardized methodologies and expanded research beyond food-related paradigms. Such advances will not only clarify the central mechanisms of GLP-1 action but also contribute to understanding how neuroendocrine signaling shapes human motivation and affect.

## Author Contributions

VD conducted the literature review and prepared the initial draft. NS substantially revised the results and discussion sections and prepared the summary figure on brain imaging. LY critically revised the manuscript. FV conceptualized the project, provided guidance during literature review, and critically revised the introduction and overall manuscript. All authors approved the final version of the manuscript.

## Acknowledgements

Research reported in this publication was supported by the National Institute on Drug Abuse of the National Institutes of Health under Award Numbers R01DA062720, R01DA053241, and R25DA059907; the MD Anderson Cancer Center Support Grant (National Cancer Institute Award Number P30CA016672); and the Baylor College of Medicine Houston Nutrition Obesity Research Center. The content is solely the responsibility of the authors and does not necessarily represent the official views of the National Institutes of Health.

